# Sexual dimorphism in the *Drosophila* metabolome increases throughout development

**DOI:** 10.1101/073148

**Authors:** FC Ingleby, Edward H Morrow

**Affiliations:** Evolution, Behaviour and Environment Group, School of Life Sciences, University 7 of Sussex, John Maynard Smith Building, Falmer, Brighton, UK

**Keywords:** *Drosophila melanogaster*, metabolome, ontogeny, sexual dimorphism, transcriptome

## Abstract

The expression of sexually dimorphic phenotypes from a shared genome between males and females is a longstanding puzzle in evolutionary biology. Increasingly, research has made use of transcriptomic technology to examine the molecular basis of sexual dimorphism through gene expression studies, but even this level of detail misses the metabolic processes that ultimately link gene expression with the whole organism phenotype. We use metabolic profiling in Drosophila melanogaster to complete this missing step, with a view to examining variation in male and female metabolic profiles, or metabolomes, throughout development. We show that the metabolome varies considerably throughout larval, pupal and adult stages. We also find significant sexual dimorphism in the metabolome, although only in pupae and adults, and the extent of dimorphism tends to increase throughout development. We compare this to transcriptomic data from the same population and find that the general pattern of increasing sex differences throughout development is mirrored in RNA expression. We discuss our results in terms of the usefulness of metabolic profiling in linking genotype and phenotype to more fully understand the basis of sexually dimorphic phenotypes.

## Introduction

Sexual dimorphism is common across a wide range of plant and animal species, and there is a longstanding field of research examining how the sexes can differ so markedly when they share the majority of their genes (Darwin 1871; Lande 1980). Most recently, research has built on the premise that for sexually dimorphic phenotypes to develop from the same genes, there is likely to be sex differences at the molecular level (Ellegren and Parsch 2007). As such, a proliferation of data on sex-biased gene expression has shed some light on the molecular basis for sex differences. Sex-biased gene expression has been found in a diverse range of species - but especially in model insect species where high throughput -omic technologies are increasingly cheap and available - and the extent of sexual dimorphism at the level of the transcriptome can be large (reviewed by Ingleby et al. 2015).

Some research has also examined how sex differences in gene expression progress throughout development. At a phenotypic level, it is generally the case that sexual dimorphism increases throughout development, ultimately resulting in highly dimorphic adult phenotypes that are adapted to sex-specific reproductive roles. Research has shown that this progression through development is often mirrored by transcriptomic sex differences. In *Drosophila melanogaster*, for instance, sex-biased gene expression is more prevalent in adults than in pre-adult stages, as demonstrated by Ingleby et al. (2016) and is also clear from comparisons of pre-adult data (e.g. Perry et al. 2014) with adults (e.g. Innocenti and Morrow 2010). Similarly, the extent of sex-biased gene expression has been shown to increase throughout development in the mosquito, *Anopheles gambiae* (Magnusson et al. 2011), and the silk moth, *Bombyx mori* (Zhao et al. 2011).

However, there is a considerable gap between gene expression and the whole organism phenotype, with a series of cellular and metabolic processes linking gene to phenotype. This may be particularly relevant in a developmental context, where the expression of a gene at a particular stage could act to trigger a pathway where the phenotypic effect might only be measurable at a later stage. This highlights the potential significance of the processes that link the genotype and phenotype. Here, we examine this gap by quantifying the metabolic profile, or metabolome, of male and female *D. melanogaster* at three stages throughout development (i.e. larvae, pupae and adults).

The usefulness of this approach is highlighted by research that illustrates the sizeable gap between genotype and phenotype - generally only a small fraction of phenotypic variation is thought to be explained by genetic variation, whereas over 50% of metabolic variation can be explained by genetic variation (Suhre et al. 2011). *Drosophila melanogaster* has been cited as a particularly useful insect model for metabolomic studies (e.g. Chintapalli et al. 2013), and recent studies have identified many interesting patterns of metabolome variation in this species: for example, variation across the sexes (Hoffmann et al. 2014) and with age (Sarup et al. 2012, Hoffmann et al. 2014), as well as metabolic plasticity to various environmental factors as adults (Overgaard et al 2007, Colinet et al. 2012, Laye et al. 2015, Williams et al. 2015) and as larvae (Kostal et al. 2011).

We used GCMS analysis to identify and quantify compounds in the *D. melanogaster* metabolic profile from male and female samples of larvae, pupae and adults. Our results show that there is significant metabolome variation both throughout development and across sexes. In addition, we find that the extent of sexual dimorphism in the overall metabolic profile increases throughout development, and that this pattern is broadly mirrored in transcriptomic data from previous research. We discuss these results in terms of how metabolic profiling could be a useful tool for further research linking genotype to phenotype.

## 2. Materials and methods

### 2.1 Fly stocks

*Drosophila melanogaster* samples for metabolic profiling were sampled from the established ‘LH_M_’ population that has been reared in consistent laboratory conditions for more than 500 generations. This population has been maintained as a large outbred population with overlapping generations, using a standard molasses diet at 25°C, 65% relative humidity, and a 12:12h light:dark incubator light cycle.

### 2.2 Sample collection and processing

Male and female flies from the stock population were given 48h to interact and mate, before males were removed, and females transferred to fresh vials of lightly yeasted food to lay eggs. Females laid eggs in these vials for 2h before being transferred to fresh vials, and then there were two subsequent 2h laying periods in fresh vials at 4 and 7 days later. This process created 3 sets of staged vials with developing offspring. For each set of vials, larvae were sexed at days after laying by visual inspection under a dissecting microscope. At this point, developing testes can be clearly seen through the larval body wall of males, allowing male and female offspring to be separated into sex-specific vials to continue development. Ten larvae were counted per vial to standardise the rearing environment. Samples were collected 11 days after the initial laying vials were set up. At this point, third instar larvae, pupae and 1-day old virgin adults (unable to mate as they eclosed in sex-specific vials) were collected from each vial. Three individuals were pooled into an eppendorf vial from each of 4 replicate vials per developmental stage and sex (N = 24 independent replicates split equally across 2 sexes x 3 developmental stages).

All samples were immediately flash frozen in liquid nitrogen, then processed following Hoffmann et al. (2014) by thoroughly homogenising the sample with a pestle motor in 150μl acetonitrile in water (2:1 v/v). Samples were then centrifuged at 12,000rpm for 20 minutes, and the 100μl of supernatant pipetted into a fresh eppendorf. Samples were stored at -80°C before being analysed approximately one week later. This involved loading the samples randomly into a chilled autosampler, and injecting 20μl of sample into a GCMS (Agilent 133 6890/5973) fitted with a DB-5MSUI column of 30m × 0.25 internal diameter × 0.25μm film thickness. Hydrogen was used as a carrier gas. The inlet was set at 280°C and the injection was in split mode. Separation of the extract was optimised with a temperature cycle that held at 50°C for 1 min, then increased at 10°C min^-^1 to 320°C. Integration of metabolite peaks was carried out using GC ChemStation software (Agilent version B.04.02.SP1), but a clear signal could not be detected for one male adult replicate, so the full analysis comprises N = 23 samples in total. Across all samples, 25 peaks were quantified and identified using mass spectroscopy data in the AMDIS software v.2.71 (Table 1). Of these compounds, 14 were present in both sexes and all stages. This indicates a considerable degree of qualitative variation in metabolome throughout development, but as our analyses focus on quantitative variation, the analysis uses only the 14 common peaks as identified in Table 1.

### 2.3 Data handling and analysis

All analyses were carried out in R v.3.2.1. Data from the integrated peaks of all 25 compounds listed in Table 1 were used to calculate standardised peak areas using a centred log ratio transformation on proportional peak areas (Pawlowsky-Glahn and Buccianti 2011), as follows:

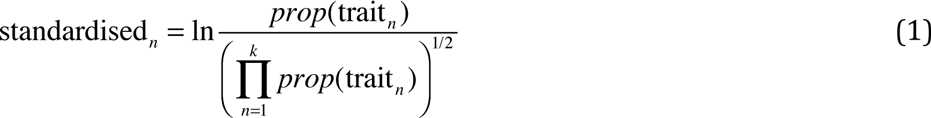

where the divisor is the geometric mean of the proportional area of all *k* traits and the numerator is the proportional area of the *n*^th^ trait. By using all peaks for the standardisation calculation, and then filtering data afterwards to leave only the 14 compounds expressed in both sexes and all stages, this avoids the problem of the zero-sum constraint of full rank data using this transformation. In total, data on expression of 14 compounds was used in further analysis in order to examine quantitative variation in metabolic profile across sexes and development.

**Table 1.** Full list of compounds identified via GCMS. The last three columns indicate whether the compound was present in each developmental stage, where ‘m’ indicates presence in male samples, ‘f’ indicates presence in female samples, and ‘fm’ indicates presence in both sexes. There is clear qualitative variation in metabolic profile throughout development. The analyses here aimed to examine quantitative variation only, and therefore used data only from compounds found in both sexes in all three developmental stages (shaded).

| Compound | Group | Stage |  |  |
| --- | --- | --- | --- | --- |
|  |  | Larvae | Pupae | Adults |
| Lactic acid | intermediate | m | fm | fm |
| Alanine | amino acid | fm | fm | fm |
| Valine | amino acid | fm | fm | fm |
| Glycerine | intermediate | fm | fm | fm |
| Leucine | amino acid | - | fm | fm |
| Glycine | amino acid | - | fm | fm |
| beta-Alanine | amino acid | - | - | fm |
| Pyroglutamic acid | intermediate | - | fm | fm |
| Glutamic acid | amino acid | - | fm | - |
| Citric acid | intermediate | fm | fm | fm |
| Inositol | polyol | fm | fm | fm |
| Fructose | sugar | - | fm | - |
| Methyl-malonic acid | intermediate | - | fm | - |
| Glucose | sugar | fm | fm | fm |
| Lysine | amino acid | - | - | fm |
| Ribonic acid | intermediate | - | fm | - |
| Palmitic acid | fatty acid | fm | fm | fm |
| Uric acid | intermediate | - | fm | - |
| Butyl palmitate | fatty acid | fm | fm | fm |
| Linoleic acid | fatty acid | fm | fm | fm |
| Oleic acid | fatty acid | fm | fm | fm |
| Stearic acid | fatty acid | fm | fm | fm |
| Butyl stearate | fatty acid | fm | fm | fm |
| Undecane | hydrocarbon | fm | fm | fm |
| Cholesterol | intermediate | fm | fm | fm |

The analyses employ a combination of univariate and multivariate approaches in order to examine variation in metabolism both as an overall metabolic profile as well as for individual compounds. Exploratory initial analyses involved hierarchical clustering of the samples based on a distance matrix using ‘hclust’ and ‘dist’ functions in the R package ‘stats’. The same clustering methods were also carried out for the 14 metabolite compounds.

Next, univariate linear models were used to directly test for sex and stage variation in expression of each of the 14 compounds individually. These linear models used Bayesian inference within the ‘MCMCglmm’ package v2.22.1 (Hadfield 2010) and each took the basic structure:

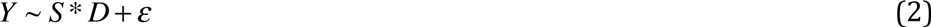

where the response variable, *Y*, represents the standardised peak area for a given compound, *S* is a fixed 2-level factor defining sex, *D* is a fixed 3-level factor representing developmental stage, and ∊ accounts for residual error variation. All models assumed a normal distribution and this assumption was checked for all compounds, as were model checks for Markov chain mixing and autocorrelation. Models used a flat prior distribution and were ran for 100,000 iterations, with a 10,000 burn-in and a thinning interval of 25. Significant differences in compound expression were inferred where 95% credible interval estimates from the posterior distribution for each sex and stage combination were non-overlapping. Note that although the main results shown are based on these Bayesian analyses, the equivalent frequentist linear models produce qualitatively identical results (Table S1).

Sexual dimorphism in the overall metabolic profile was examined via linear discriminant analysis, where differentiation between male and female samples was modelled as a function of all 14 compounds. From the results of this model, each sample was given a score along the discriminant function vector defining maleness/femaleness, and these scores were modelled using the approach described for equation [2], where *Y* in this instance is the discriminant function score for each sample.

Finally, variation in metabolic profile across sexes and development was compared to variation in gene expression found across the same sample types, derived from the same *D. melanogaster* population, in terms of gene expression. This data is taken from a previous study of the population, where RNA-204 sequencing was carried out on male and female samples of larvae, pupae and adults (Ingleby et al. 2016). Here, we filtered the transcriptome to focus on a subset of 26 genes that were identified as involved with the tricarboxylic acid (TCA) cycle, using the database of *D. melanogaster* genes in the R package ‘biomaRt’ (filtered using ‘grep’ for the term ‘TCA’ in the gene description field). This filter was applied based on the results of the metabolite analysis, which showed interesting patterns of expression for compounds involved in the tricarboxylic acid cycle. After filtering the RNA expression data to only include these genes, a linear discriminant analysis was carried out exactly as described for the metabolite data, and the resulting scores were modelled as before.

## 3. Results

Initial hierarchical clustering of samples indicated strong differentiation between developmental stages, with all samples grouped by stage (Figure 1). Larval and pupal branches were more closely linked, with the adult branch as an out-group. Within each stage, male and female samples did not cluster separately, although arguably there was more evidence of sex differentiation in adults than in the earlier developmental stages, since the adult male samples clustered together (Figure 1).

**Figure 1.**
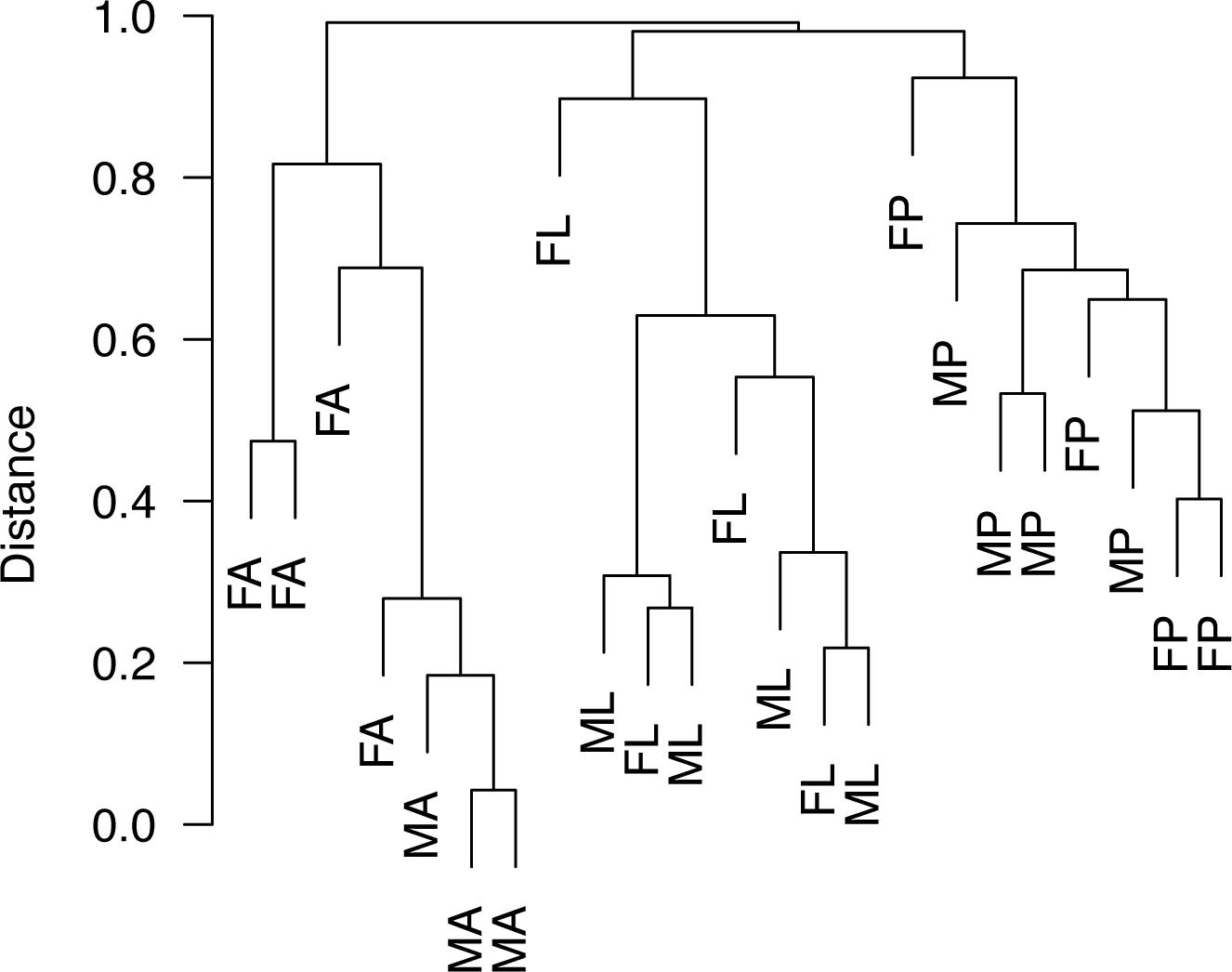
Result of hierarchical distance clustering to differentiate between samples. The distance is shown on the y-axis scale, and length of branches corresponds to the distance. The tree is based on expression of 14 compounds that were present at some level in all samples.

Cluster analysis of the 14 metabolites that were found in all sample types (as described in Table 1) showed a tendency for fatty acids and fatty acid derivatives to group together (Figure 2), suggesting that these compounds were expressed more similarly to each other than the other compounds analysed (predominantly sugars and amino acids).

**Figure 2.**
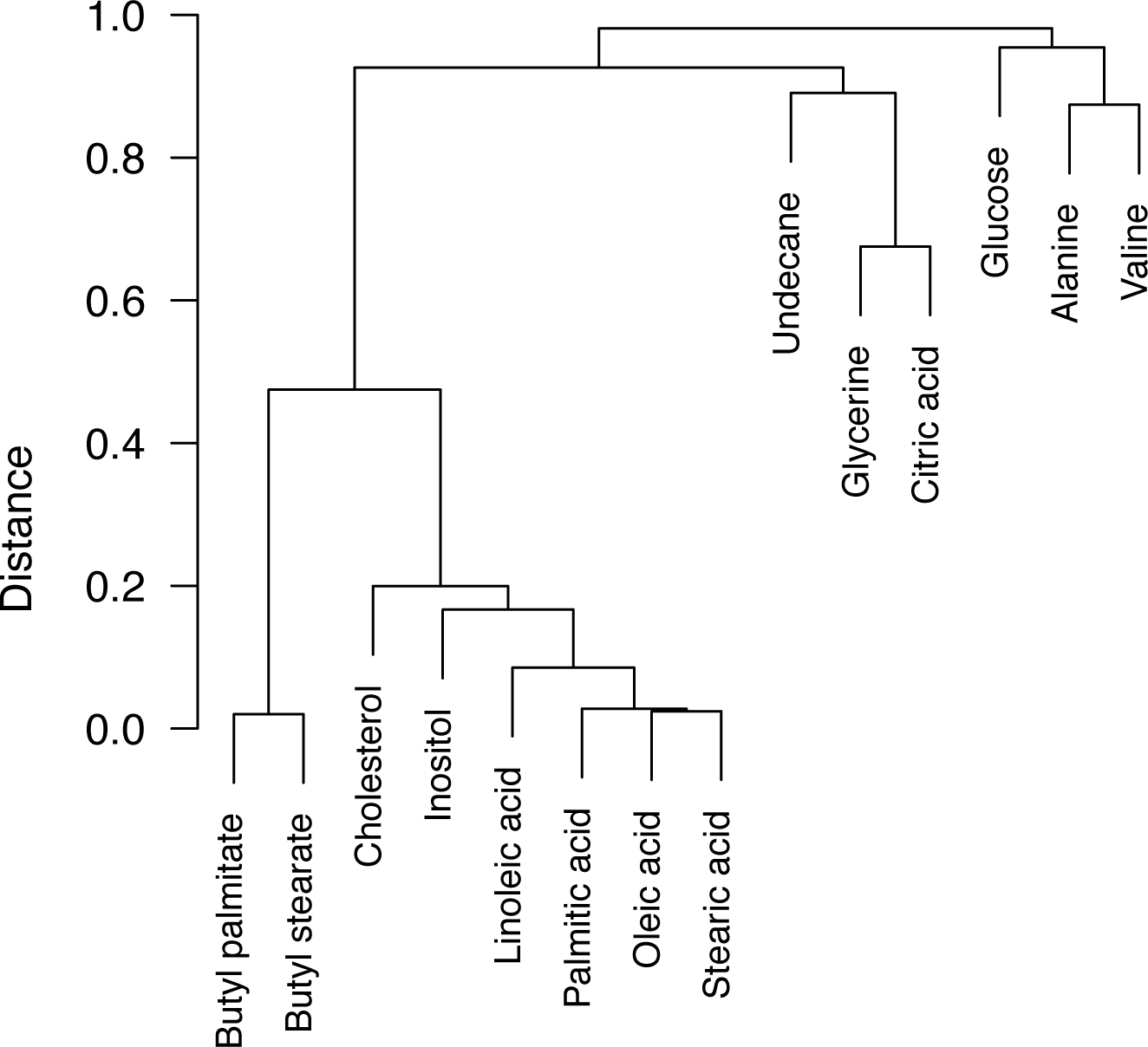
Result of hierarchical distance clustering to differentiate between metabolite compounds. The distance is shown on the y-axis scale, and length of branches corresponds to the distance value. All compounds were present at some level in all samples included in this analysis.

Significant differences in the expression of individual compounds across sexes and developmental stages were tested directly via linear model analysis. Highly significant differentiation across stages was found for glycerine, citric acid, glucose and undecane (Figure 3), with further evidence of a significant sex x stage interaction in the expression of undecane. Visual inspection of the posterior distribution estimates in Figure 3 also suggests a possible sex x stage interaction for alanine and citric acid expression, but overlap between the 95% credible intervals for the different sample types shows that these interactions are non-significant. Note that it is likely that relatively small sample sizes have contributed to wide credible intervals (indicating wide variance around the posterior estimates).

**Figure 3.**
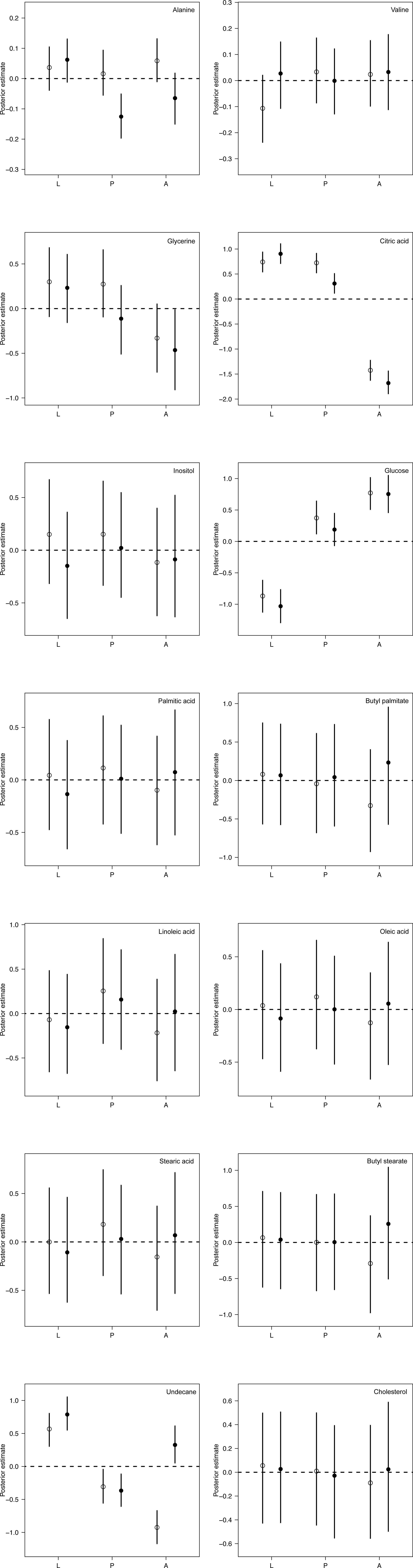
Posterior estimates (mean with 95% credible intervals) for each sex and stage combination taken from univariate MCMCglmm linear models for each of the 14 compounds (named in the top right of each plot). Female estimates are shown with open points, male estimates with filled points. L = larvae; P = pupae; and A = adults.

Despite non-significant sex differences in the univariate analyses, multivariate analyses provide convincing evidence for sex differentiation between males and females of the metabolic profile overall. A linear discriminant analysis clearly differentiated between the sexes. This model was used to project samples onto the linear discriminant vector LD1, giving each sample a score along an axis describing metabolomic sex differentiation. These scores were significantly different between male and female pupae and adults, although not between male and female larvae (Figure 4). In addition, the difference between male and female posterior means tends to increase throughout development (absolute difference in posterior mean male and female scores in larvae = 1.34; pupae = 3.01; and adults = 3.85), suggesting an increase in the extent of metabolome sexual dimorphism from larvae through to adults.

**Figure 4.**
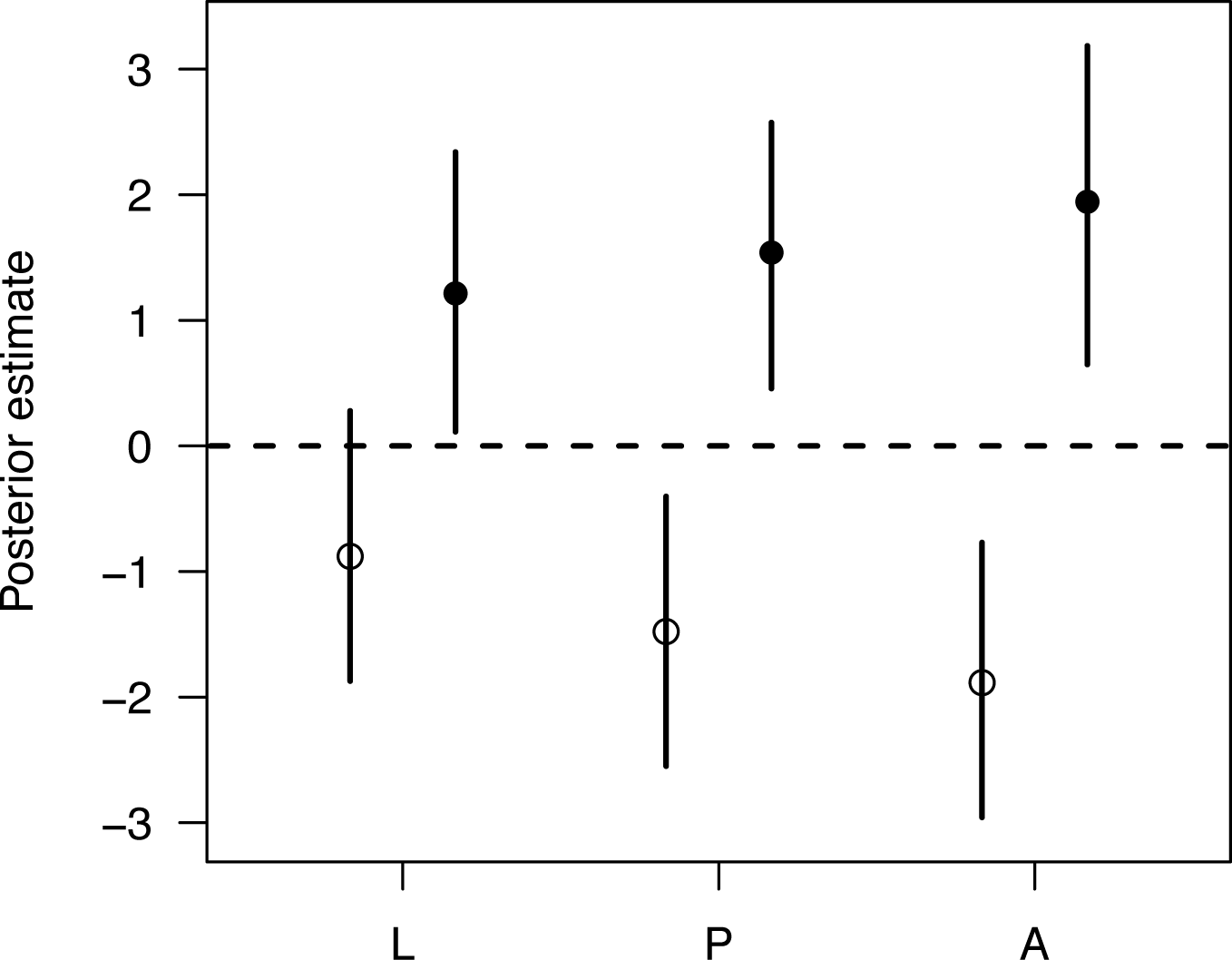
Posterior estimates (mean with 95% credible intervals) for females (open points) and males (filled points) from each developmental stage for the first linear discriminant function (LD1). These values represent scores along a discriminant vector that differentiates between the sexes.

The equivalent multivariate analysis based on gene expression data of genes associated with the tricarboxylic acid cycle (see Methods) revealed a very similar pattern in sexual dimorphism of the transcriptome throughout development. Scores for LD1 in this case were significantly different between males and females at all three developmental stages (Figure 5), and the difference between male and female posterior mean estimates tends to increase throughout development (absolute difference in posterior mean male and female scores in larvae = 4.73; pupae = 5.43; and adults = 6.41).

**Figure 5.**
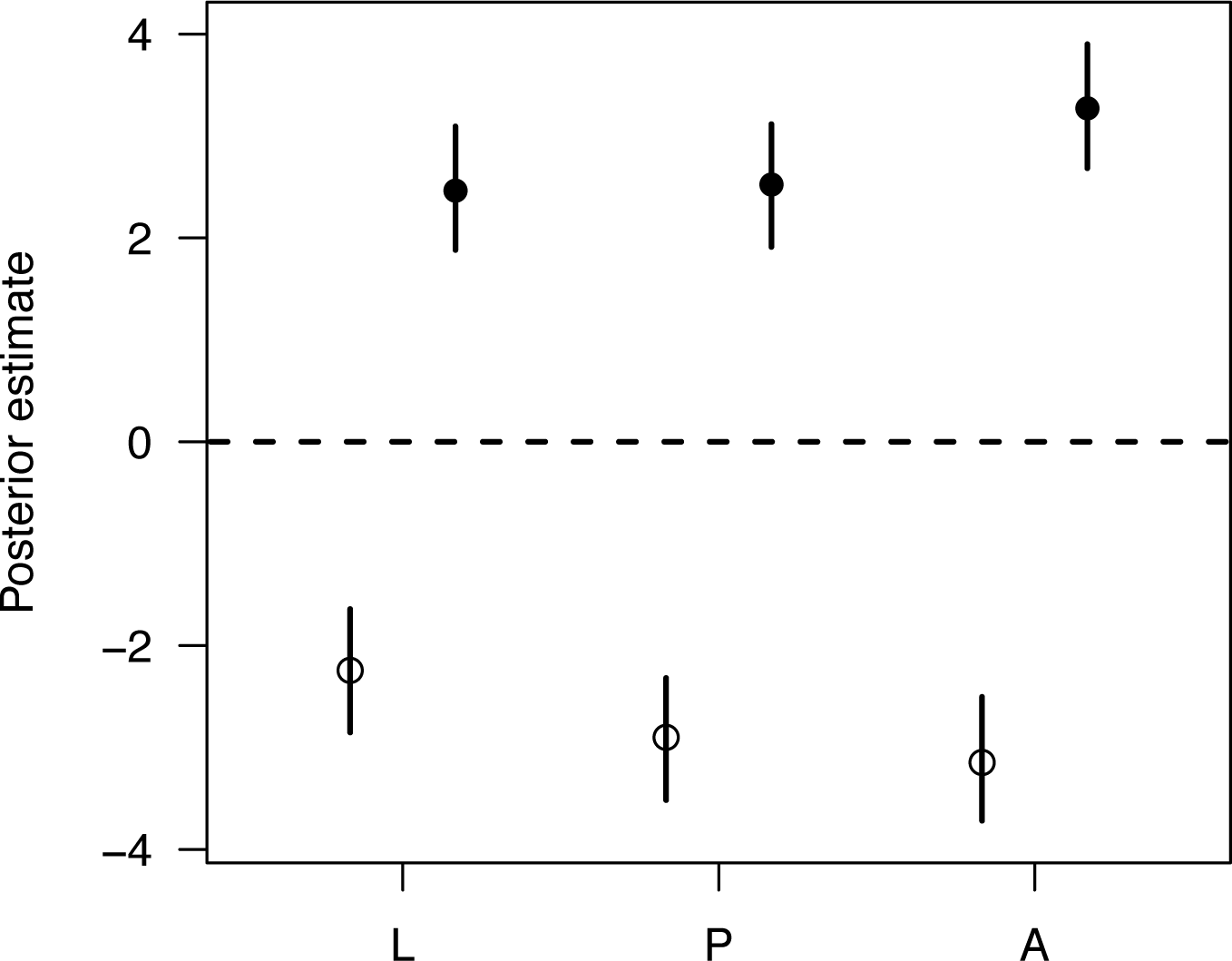
Posterior estimates (mean with 95% credible intervals) for females (open points) and males (filled points) from each developmental stage for the first linear discriminant function (LD1) based on RNA data from 26 genes identified as associated with the tricarboxylic acid cycle. These values represent scores along a discriminant vector that differentiates between the sexes.

## 4. Discussion

Recent research has found variation in the *D. melanogaster* metabolome across different environments, ages, and sexes (e.g. Colinet et al. 2012, Hoffmann et al. 2014, Laye et al. 2015, Williams et al. 2015). In this study, we found clear variation in metabolic profile across larval, pupal and adult developmental stages, as well as sex differences in the overall profile that appear to increase throughout development. Additionally, we present some evidence that suggests sex differences in the metabolome throughout development are mirrored by ontogenetic patterns in the sex-biased expression of related genes. Our analyses focussed on the evidence for quantitative variation in metabolites, but we also found considerable qualitative variation throughout development, since almost half of the individual metabolites identified in our samples were stage- or sex specific. This qualitative variation is not examined beyond identification here, but may be of interest for future research.

Evidence for quantitative metabolome variation across development was very clear both from the multivariate analysis of the overall metabolic profile, as well as from univariate analysis of individual metabolites. Three of the metabolites that varied significantly between developmental stages - glucose, citric acid and glycerine - are key components in the tricarboxylic acid cycle, or Krebs cycle, that is largely responsible for providing cells with energy (Baldwin and Krebs 1981). Generally, our data shows an increase in glucose throughout development, combined with a decrease in citric acid and glycerine. While these patterns are interesting, it is difficult to disentangle any functional significance without a more detailed dataset, so here we simply note that the significant variation in these chemical components indicates that the dynamics of this cycle may vary throughout development. More generally, developmental variation in metabolism has been found previously related to diapause in various insect species (Hahn and Denlinger 2007, Michaud and Denlinger 2007, Li et al. 2015, Dean et al. 2016), and Callier et al. (2015) found that metabolic responses to anoxia differed between larvae and adults in *D. melanogaster*.

The strong differences between stages of these specific metabolites directed our analysis of the transcriptome data, which focussed on a subset of 26 genes that were associated with the cell tricarboxylic acid cycle. In fact, from the gene descriptions in the ‘biomaRt’ database, many of these genes code for dehydrogenase enzymes directly involved with different steps of the cycle. This analysis showed an increasing extent of sexual dimorphism in gene expression through development, and this supports the idea that these metabolites and the associated genes could be interesting candidates for further research into sex and stage differences in the metabolome.

The patterns of sex dimorphism in the RNA data mirrored those that we found from the multivariate analysis of the overall metabolomic profile, although none of the individual metabolites tested significantly for sex dimorphism in the univariate analyses. In part, these non-significant results could be due to a relatively small sample size, which would mean that power to detect differences would be low, and this is supported by the wide intervals on the posterior estimates. Indeed, sex differences in various aspects of metabolism have been identified in previous studies of animals as diverse as *D. melanogaster*(Hoffmann et al. 2014) and humans (Kochhar et al. 2006), and so the lack of significant sex dimorphism for individual metabolites here is unexpected. However, the multivariate profiling approach revealed sexual dimorphism in the overall metabolome in the later stages of development (pupae and adults), and a general trend for the difference between the male and female metabolome to steadily increase throughout development. This increase in the extent of sexual dimorphism largely reflects the broad pattern of phenotypic sexual dimorphism increasing throughout development - phenotypically, males and females diverge throughout development, ultimately resulting in dramatically different adult phenotypes that are well-adapted to sex-specific roles (Darwin 1871).

This pattern of sexual dimorphism throughout development mirrors the results of our transcriptomic analyses and is also consistent with other studies that show an increasingly sexually dimorphic transcriptome throughout development (Magnusson et al. 2011, Zhao et al. 2011, Ingleby et al. 2016). Our attempt to link previous transcriptomic data with the new metabolomic data was intended as exploratory only. As such, further research could undoubtedly improve on this straightforward comparison of two datatsets. More generally, the attempt to link metabolomic and transcriptomic data highlights the availability of detailed -omic data for *D. melanogaster*, and although to a lesser extent, other insect model species as well. The increasing use of genomic, transciptomic and metabolomic technology should mean that we can reach closer to a full understanding of how genotype maps to phenotype - including the oft-overlooked steps between the gene and the whole organism phenotype. This will be instrumental in future sexual dimorphism research, since one of the key problems that a genotype phenotype map could address is how different phenotypes (i.e. sexes) are produced from the same genes. With regards to the metabolome, this is unlikely to be a simple linear map from gene to metabolite to phenotype. Recent research has indicated, for instance, that environmental influences on the metabolome could form a basis for interaction effects between genotype and phenotype (Williams et al. 2015). Although complex, this suggests that more in-depth metabolomic profiling - for example, within a quantitative genetic framework could be useful in understanding how different phenotypes are formed across different contexts, including different sexes and developmental stages of development.

## Acknowledgements

The authors are grateful to the Mass Spectrometry Facility at King’s College London for their help processing the samples, and to Chris Mitchell for his help with using the ChemStation software. This work was supported by funding from the Swedish Research Council (2011-3701), the European Research Council (#280632), and a Royal Society University Research Fellowship (all to E.H.M).

---

**Table S1.** Results of non-Bayesian univariate linear models testing for differences in metabolite expression between developmental stages, sexes, and the interaction. For each effect, the F statistic and associated P value are shown. P values are corrected for FDR < 0.05 and significant results are highlighted in bold. Models (described in the text) were equivalent to the Bayesian univariate linear models, and all results are qualitatively the same through Bayesian inference (shown in main text). Note that the interaction effect was significant for alanine and citric acid prior to FDR correction.

| Compound | Stage x sex |  | Stage |  | Sex |  |
| --- | --- | --- | --- | --- | --- | --- |
|  | F | P | F | P | F | P |
| Alanine | 3.45 | 0.25 | 3.60 | 0.13 | 5.81 | 0.18 |
| Valine | 1.04 | 0.87 | 0.74 | 0.96 | 0.57 | 0.83 |
| Glycerine | 0.43 | 0.87 | **7.07** | **0.02** | 1.90 | 0.60 |
| Citric acid | 5.09 | 0.13 | **236.90** | **< 0.001** | 3.03 | 0.46 |
| Inositol | 0.22 | 0.87 | 0.36 | 0.96 | 0.53 | 0.83 |
| Glucose | 0.25 | 0.87 | **105.58** | **< 0.001** | 1.66 | 0.60 |
| Palmitic acid | 0.27 | 0.87 | 0.12 | 0.96 | 0.05 | 0.99 |
| Butyl palmitate | 0.42 | 0.87 | 0.12 | 0.96 | 0.57 | 0.83 |
| Linoleic acid | 0.22 | 0.87 | 1.03 | 0.88 | 0.01 | 0.99 |
| Oleic acid | 0.23 | 0.87 | 0.12 | 0.96 | 0.03 | 0.99 |
| Stearic acid | 0.26 | 0.87 | 0.28 | 0.96 | 0.01 | 0.99 |
| Butyl stearate | 0.44 | 0.87 | 0.04 | 0.96 | 0.37 | 0.85 |
| Undecane | **15.06** | **0.002** | **19.35** | **< 0.001** | 7.43 | 0.18 |
| Cholesterol | 0.07 | 0.93 | 0.07 | 0.96 | 0.01 | 0.99 |

